# Functional Bias and Demographic History Obscure Patterns of Selection among Single-Copy Genes in a Fungal Species Complex

**DOI:** 10.1101/107326

**Authors:** Santiago Sánchez-Ramírez, Jean-Marc Moncalvo

## Abstract

Many different evolutionary processes may be responsible for explaining natural variation within genomes, some of which include natural selection at the molecular level and changes in population size. Fungi are highly adaptable organisms, and their relatively small genomes and short generation times make them pliable for evolutionary genomic studies. However, adaptation in wild populations has been relatively less documented compared to experimental or clinical studies. Here, we analyzed DNA sequences from 502 putative single-copy orthologous genes in 63 samples that represent seven recently diverged North American *Amanita* (*jacksonii*-complex) lineages. For each gene and each species, we measured the genealogical sorting index (*gsi*) and infinite-site-based summary statistics, such as 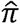, 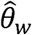 and *D_Taj_* in coding and intron regions. MKT-based approaches and likelihood-ratio-test *K_n_/K_s_* models were used to measure natural selection in all coding sequences. Multi-locus (Extended) Bayesian Skyline Plots (eBSP) were used to model intraspecific demographic changes through time based on unlinked, putative neutral regions (introns). Most genes show evidence of long-term purifying selection, likely reflecting a functional bias implicit in single-copy genes. We find that two species have strongly negatively skewed Tajima’s *D*, while three other have a positive skew, corresponding well with patterns of demographic expansion and contraction. Standard MKT analyses resulted in a high incidence of near-zero α with a tendency towards negative values. In contrast, α estimates based on the distribution of fitness effects (DFE), which accounts for demographic effects and slightly deleterious mutations, suggest a higher proportion of sites fixed by positive selection. The difference was more notorious in species with expansion signatures or with historically low population sizes, evidencing the concealing effects of specific demographic histories. Finally, we attempt to mitigate Gene Ontology term overrepresentation, highlighting the potential adaptive or ecological roles of some genes under positive selection.

## Introduction

Understanding the nature of genomic variation can help illustrate how natural selection and demography balance out throughout the history of a species. The neutral theory of molecular evolution (Kimura 1986; Ohta 1992) postulates that most mutations contributing substantially to the genetic pool of a population will be either neutral or nearly so. Behind this reasoning is the assumption that changes that alter the amino acid composition of a protein will tend to be deleterious and selected against, while those that are advantageous will be rare. In this framework, one can test whether DNA variation at a given locus fits the expectations under the neutral equilibrium model (Tajima 1989; Fu and Li 1993, 1999). These models primarily allow for explicit testing of neutrality departures, however different kinds of evolutionary processes, such as selective sweeps, recombination, and demographic changes, can mimic similar results (Bachtrog and Andolfatto 2006; Ramírez-Soriano et al. 2008). Nonetheless, selection is likely to affect specific regions through the fixation of linked neutral variation –a process known as hitchhiking or selective sweeps–, while demographic changes are more likely to have genome-wide effects (Andolfatto 2001; Nielsen 2001).

Natural selection at the molecular level is detectable in protein-coding DNA. This can be accomplished by looking at the relationship between functional (amino acid-changing or non-synonymous) and silent (neutral or synonymous) variation. A widely use test of selection is the McDonald-Kreitman test (MKT, McDonald and Kreitman 1991), which is based on the comparison between the ratio of functional (*π*_*n*_) and silent (*π*_*s*_) mutations within a species (polymorphism) and the ratio of fixed *K*_*n*_ and *K*_*s*_ mutations between species (divergence). If both polymorphism and divergence were solely driven by mutation and drift, one would expect their ratio to equal one. This model assumes that all *K*_*s*_ mutations are neutral and that *K*_*n*_ mutations are either strongly deleterious, neutral or strongly advantageous (Smith and Eyre-Walker 2002). In this framework, advantageous mutations are more likely to contribute to divergence due to their high fixation probability, while deleterious mutations are likely to be found at lower frequencies due to the effects of purifying selection. On the other hand, the vast majority of segregating mutations are likely to be neutral, as they will tend to persist in the population for longer periods until they eventually go to fixation by random genetic drift (Sella et al. 2009). However, the proportion of adaptive mutations can be under-or overestimated under certain demographic scenarios combined with weak selection (Smith and Eyre-Walker 2002; Eyre-Walker 2006; Messer and Petrov 2013a). For instance, in species with reduced population sizes, slightly deleterious mutations are effectively neutral, thus they are at higher frequency in the population resulting in an underestimation of the proportion of adaptive mutations. Similarly, a recent population expansion will tend to decrease polymorphism, confounding sweep-driven positive selection (Nielsen 2001; Nielsen et al. 2005; Wright and Gaut 2005). Furthermore, another approach is to quantify selection directly as the ratio of substitution rates at *K_n_* and *K_s_* sites (Kimura 1977; Huges and Nei 1988; Goldman and Yang 1994). In a neutral scenario, *K_n_/K_s_* or *ω* should approximate 1, with values above indicating diversifying (positive) selection, and values below indicating purifying (negative) selection (Nielsen 1997; Yang and Nielsen 2000; Nielsen 2001).

Detecting genome-wide patterns of selection and demography is now feasible through the growing availability and accessibility of whole-genome data; the field now known as population genomics (Charlesworth 2010). Most studies addressing questions such as genome-wide effects of selection have usually focus on a number of model organisms (Hough et al. 2013; Ellegren 2014), most of which have good reference genomes and well-documented annotations, while for the vast majority of species, empirical evidence is still lacking (Cutter and Payseur 2013). Nonetheless, lowering sequencing costs and more efficient bioinformatics tools are making these types of approaches accessible to non-model organisms (Ekblom and Galindo 2011; Aguileta et al. 2010).

Fungi play multiple essential roles in the environment, mainly as decomposers of organic matter, but also as pathogens, commensals, and mutualists. Not only due to their diverse ecology (Selbmann et al. 2013), but also because of their genetic machinery (Anderson et al. 1992; Schoustra et al. 2007), fungi are regarded as highly adaptable organisms. Most evidence comes from experimental studies in yeasts (Suutari et al. 1990; Davies et al. 1995; Piper et al. 2001; Liu 2006; Dettman et al. 2007; Anderson et al. 2010; Gerstein et al. 2014), drug resistance (Kontoyiannis and Lewis 2002; Anderson 2005), and the evolution of virulence in pathogens (Bentrup and Russell 2001; Becher et al. 2010; Fisher et al. 2012). In spite of being great candidates (mostly due to relatively smaller genomes and short generation times) to study evolutionary genomics (Gladieux et al. 2014), adaptation in wild fungal populations and species has been scarcely explored. However, recent work in few fungal species has evidenced adaptive mechanisms molecularly and experimentally in the wild. For instance, in *Neurospora*, environmental factors such as temperature and latitude are contributing components of genome divergence and adaptation in wild populations (Ellison et al. 2011). Moreover, Aguileta et al. (2012) and Gladieux et al. (2013) highlight the importance of selection and molecular adaptation during host switches and host specialization in *Botrys* and *Microbotryum*. Finally, Branco et al. (2015) explores patterns of genome divergence in two populations of the ectomycorrhizal (EM) —root-associated symbionts— fungus *Suillus* adapted coastal and montane environments. These studies are few recent examples that evidence genome-wide adaptive mechanisms in wild fungal populations.

The genus *Amanita* comprises 500-1000 species (Tulloss 2005), most of which live as EM symbionts (Wolfe et al. 2012). The Caesar’s mushrooms (sect. *Caesareae*) form a clade of edible EM *Amanita* distributed worldwide, and are particularly diverse in North America (Sánchez-Ramírez et al. 2015a). With its eight to seven species, the *A. jacksonii* complex is an example of one of two continental Plio-Pleistocene radiations, leading to variable demographic histories and geographic ranges (Sánchez-Ramírez et al. 2015b; Fig. 1). Here we sought to disentangle demographic from adaptive processes in a multi-species comparative framework, mining a data set of 502 putative-single-copy genes produced by exon-target sequencing. We also explore functional enrichments of genes under selection and speculate about their role in adaptive and ecological processes.

**Figure 1.**
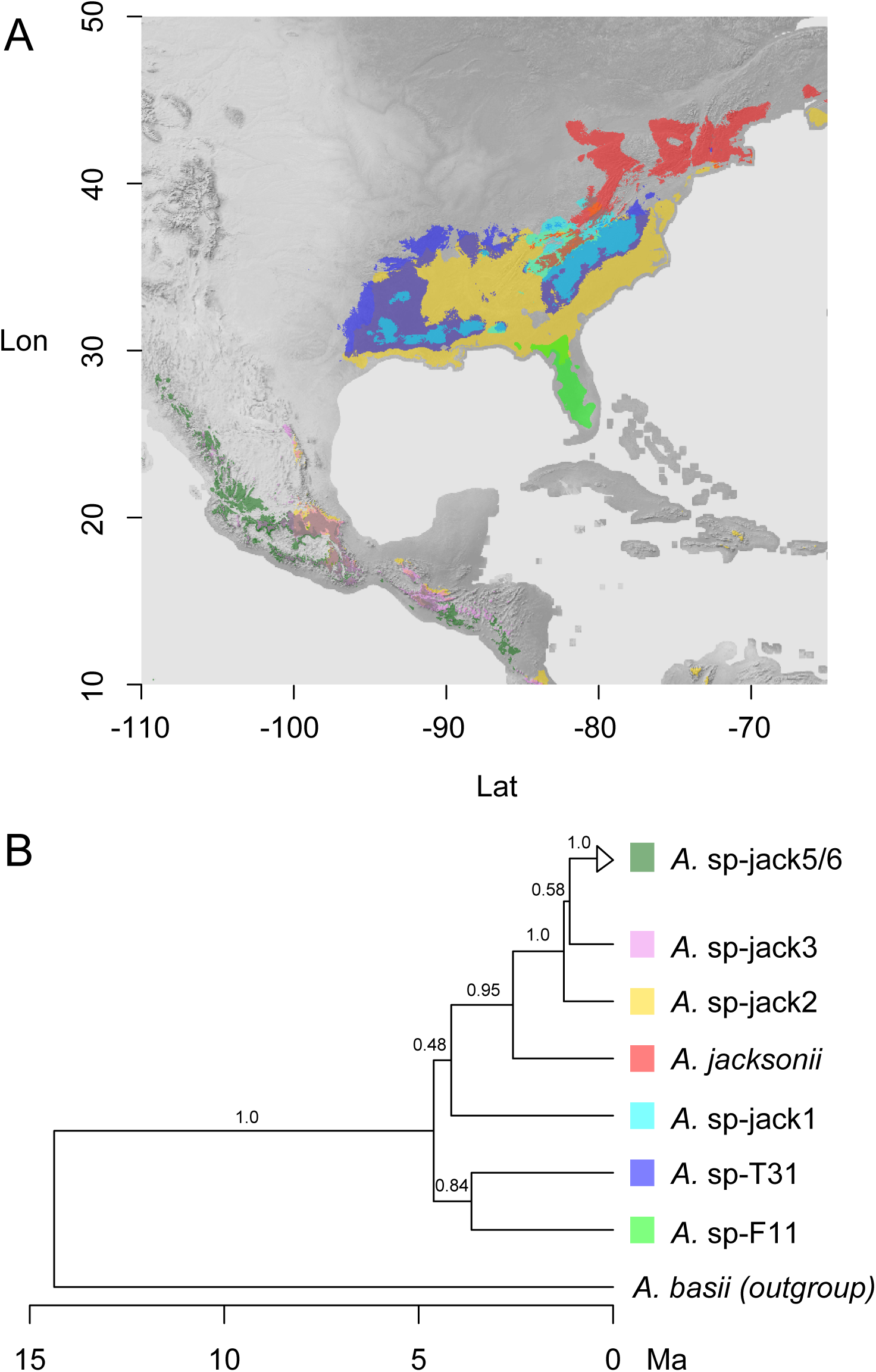
A, Species distribution maps based on geographic data from Sánchez-Ramírez et al. (2015b) and MaxEnt (Phillips et al. 2006). B, Species tree showing phylogenetic relationships and divergence times with relation to the outgroup (modified from Sánchez-Ramírez et al. 2015b). Branches show posterior probability support.

## Results

### Targets, sequence data, and bioinformatics

Reciprocal BLAST hits between predicted proteomes in the draft genomes of *A. jacksonii* and *A. basii* identified 3,427 putative single-copy candidate genes. The total number of structural genes compared was of 8,511 in *A. jacksonii* and 5,878 in *A. basii*. From the set of 3,427, we selected genes based on a criterion of a stretch of 60 bp of conserved DNA and a minimum number of four probes per gene. Most genes exceeded the minimum number criterion by having 7 to 8 non-overlapping probes. This resulted in 3,887 probes hybridizing to 502 genes.

The total number of reads per sample ranged between 1,787,146 and 24,316,811, with a mean of 9,157,472. The percentage of on-target reads per sample ranged between 56.6% and 96.3%, with an average of 81%. The average number of on-target reads per sample varied between 3,078.6 and 45,760.5 per gene, with a mean of 14,720 across samples. All 502 genes were recovered for all 63 samples, except for samples RET 109-4 and FCME12243 where one gene was not recovered, and sample FCME5528 where two genes were not recovered.

The genes ranged in size from 1,136 to 13,720 bp, with an average length of 4,950 bp. Introns were smaller in size than exons [coding DNA (CDS)], with an average of 68.56 compared to 258.69 bp in CDS. Each gene had about 14 introns and 15 CDS on average. Eighty percent of the data corresponded to CDS, while only 20% corresponded to introns. In total, the data represented 2,483,742 nucleotide sites. We measured the amount of missing data in the alignments given that low mapping quality regions and/or regions with no coverage resulted in missing data (e.g. Ns). We find that both the amount of missing data and the number of variable sites (across all 7 species) is correlated with gene size (Supplementary Fig. S1).

### Genealogical sorting and reciprocal monophyly

In order to measure how well genes supported previous species delimitations (Sánchez-Ramírez et al. 2015b) and to assess the relative contribution of phylogenetic conflict (e.g. due to introgression and/or ancestral polymorphism) to polymorphism patterns, we constructed Bayesian trees and calculated the genealogical sorting index (*gsi*, Cummings et al. 2008). The *gsi* measures the degree of co-ancestry within labeled groups and cannot distinguish between introgression and ancestral polymorphism (i.e. incomplete lineage sorting). However, it provides an overall view of how consistent gene trees are with species divergences (the index goes from 1 to 0). Measures of *gsi* indicate that the vast majority of gene trees support the monophyly of at least six species, which all have median *gsi* values above 0.8 (Fig. S2 and S3 in Online Supplementary Data). In contrast, species *A.* sp-jack5 and *A.* sp-jack6, delimited as distinct species in Sánchez-Ramírez et al. 2015b, had median *gsi* values between 0.6 and 0.4 suggesting some degree of phylogenomic conflict between them. When considered as a single unit the *gsi* value increased to 0.82 (Fig. S3 in Online Supplementary Data). For this reason, data from *A.* sp-jack5 and *A.* sp-jack6 were hereafter regarded as *A.* sp-jack5/6 and considered in the following analyses as a single species.

### DNA polymorphism

The highest 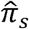 was found in *A.* sp-F11 with a mean of 0.0119, while the lowest mean, 0.0029, was found in *A.* sp-jack3 –a 3 order magnitude difference (Table 1). As expected due to selective constrain, 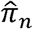 was lower in comparison with mean values ranging from 0.0005 in *A.* sp-jack3 to 0.0013 in *A.* sp-F11 and *A.* sp-jack5/6 (Table 1). The mean 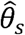 ranged from 0.0098 in *A.* sp-jack5/6 to 0.0027 in *A.* sp-jack3, while the mean 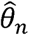 ranged from 0.0012 in *A. jacksonii* to 0.0005 in *A.* sp-jack3 (Table 1). On average, higher polymorphism was observed in synonymous sites compared to intronic sites (Table 1). We find that on average, two species (*A. jacksonii* and *A.* sp-jack5/6) had a negatively skewed Tajima’s D (*D*), in particular *A. jacksonii*; two other species (*A.* sp-jack2 and *A.* sp-jack3) had average *D* values close to 0; and three other (*A.* sp-jack1, *A.* sp-T31, and *A.* sp-F11) had positively skewed *D*_*Taj*_ values (Fig. 2a). *D*_*n*_ was on average lower than *D*_*s*_. In contrast, Fu and Way’s *H* (Fu and Way 2000), which measures the skewness of the derived (polarized by ourgroup) site-frequency spectrum (SFS), was lower in synonymous sites than in non-synonymous sites (Table 1). Due to potential bias in frequency-based estimators of polymorphism, such as Tajima’s *D*, caused by ancestral polymorphism and/or gene flow (Städler et al. 2009; Cutter and Choi 2010; Cutter et al. 2012), we looked at the relationship between *gsi* values and *D*. With the expectation that a strong effect would show a linear relationship between low *gsi* values and *D*, we found that genes with significant amounts of within-species phylogenetic conflict do not skew estimates compared to those that fully support monophyly within species (*gsi* = 1; Fig. S4 Online Supplementary Data).

**Figure 2.**
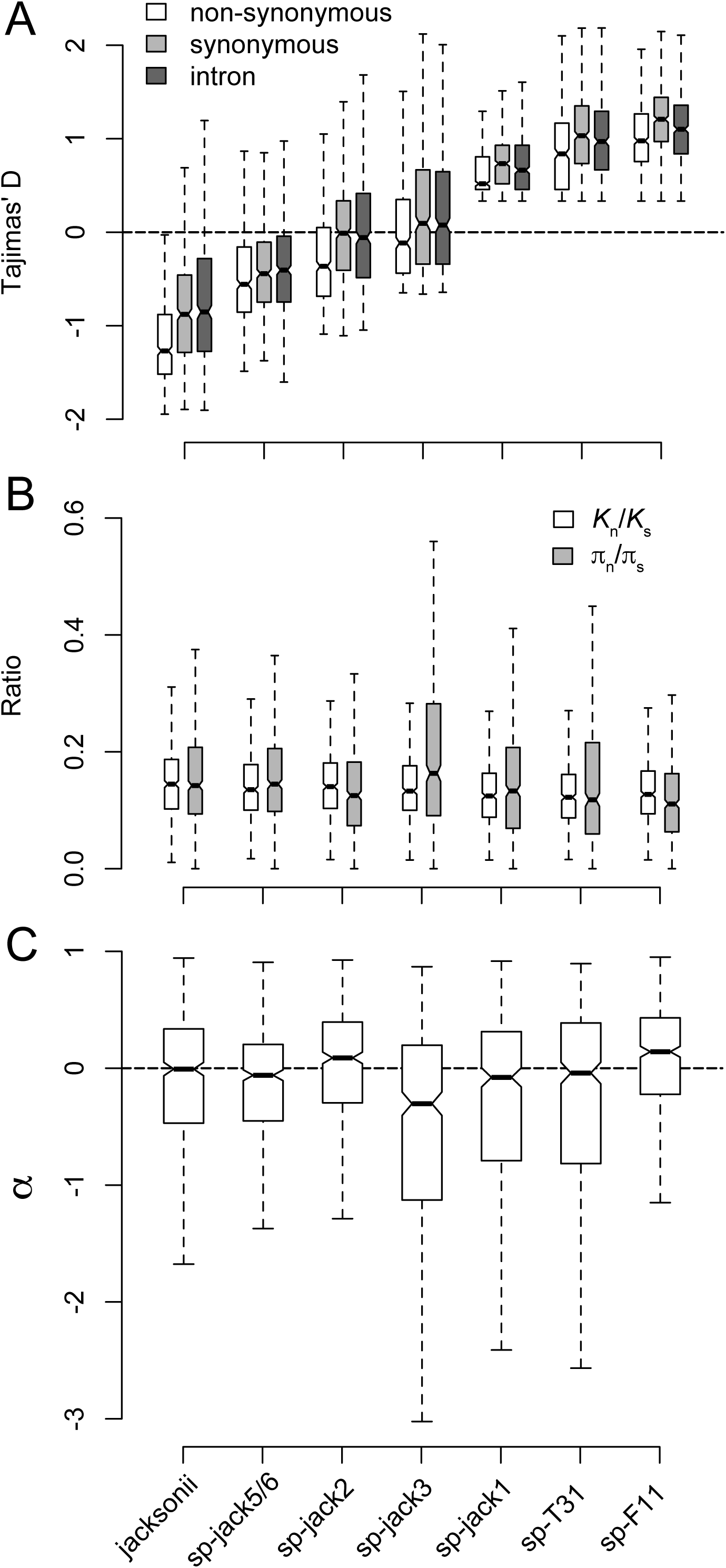
Polymorphism and divergence data for 502 genes. A, box-plots of Tajima’s D separated by site-classes (non-synonymous, white; synonymous, light grey; introns, dark grey) for the A. jacksonii complex. B, box-plots of the ratios of non-synonymous to synonymous polymorphism and divergence. C, α values from the standard MKT test.

### Divergence and rates of adaptive evolution

Divergence (*K*) consistently varied between species in the order of 10% for synonymous sites and 2% for non-synonymous sites (Table 1). The ratios between 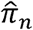 and 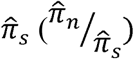 and *K_n_* and 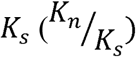 were quite close to each other, both within and between species (Fig. 2b). This resulted in neutrality index (NI) values that ranged on average from 0.977 in *A.* sp-F11 to 1.614 in *A.* sp-jack3, and *α* values close to zero with a tendency of negative values (Fig. 2c). Negative *α* values can be caused by strong purifying selection, resulting in 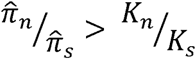as deleterious variant will not contribute to divergence, but will be found at lower frequencies as polymorphism (Nielsen 2001; Egea et al. 2008). To explore this possibility, we performed likelihood-ratio-test (LRT) of codon-based substitution models (Yang and Nielsen 2002) between species and found that most genes (278, 55%) fitted a model that supports purifying selection (the nearly-neutral model, M1a, with parameters *ω* = 1 and *ω* < 1), while only 10% (48) were supported by a diversifying selection model (the positive selection model, M2a, with parameters *ω* = 1, *ω* < 1, and *ω* > 1). For genes fitting the nearly-neutral and diversifying selection models, the mean *ω* across sites was 0.23 ± 0.09 and 0.27 ± 0.12, respectively. The rest (176, 35%) did not significantly (0.05 significance level) fit either model, so were deemed to be under stronger purifying selection, given that *ω* << 1(mean *ω* 0.18 ± 0.1). Moreover, genes with longer than expected coalescence times (i.e. ancestral polymorphism or introgression) can also lead to a 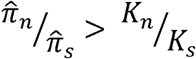 effect by increasing the frequency of weakly deleterious variants, or variants under balancing selection (Charlesworth 2009), causing negative *α* values. However, we find no relationship between *gsi* and *α* (Fig. S5 Online Supplementary Data). A third possibility is that weak selection at synonymous coding (e.g. codon usage bias) is reducing synonymous relative to non-synonymous diversity, however we find no such pattern (Table 1 and Fig. S6 Online Supplementary Data).

**Table 1.** Polymorphism, divergence, and codon usage bias estimations for 502 genes in the *A. jacksonii* complex. N, sample size; *S*, segregating sites; π, nucleotide diversity; θ, Watterson’s theta; *H*, Fay and Wu’s *H*; NI, neutrality index; CAI, codon adaptation index. Sub-indices ‘n’, ‘s’, and ‘int’ represent non-synonymous, synonymous, and intronic sites.

|  | *A. jacksonii* | *A. sp-jack56* | *A. sp-jack2* | *A. sp-jack3* | *A. sp-jack1* | *A. sp-T31* | *A. sp-F11* |
| --- | --- | --- | --- | --- | --- | --- | --- |
| N | 44 | 26 | 18 | 14 | 8 | 8 | 8 |
| *S<sub>n</sub>* | 16 (3–39) | 17 (4–39) | 12 (1–29) | 5 (0–14) | 5 (0–12) | 4 (0–11) | 9 (1–23) |
| *S<sub>s</sub>* | 25 (6–62) | 29 (12–52) | 22 (5–52) | 7 (1–20) | 8 (1–18) | 7 (1–18) | 19 (4–42) |
| *S<sub>int</sub>* | 17 (2–48) | 19 (3–45) | 15 (1–41) | 5 (0–15) | 5 (0–15) | 5 (0–14) | 13 (1–35) |
| $\hat{\pi}_n$ | 0.0008<br>(0.0001–0.0021) | 0.0013<br>(0–0.0025) | 0.001<br>(0–0.0024) | 0.0005<br>(0.0002–0.0013) | 0.0007<br>(0.0002–0.0015) | 0.0006<br>(0.0003–0.0017) | 0.0013<br>(0.0001–0.0032) |
| $\hat{\pi}_s$ | 0.0059<br>(0.0013–0.0157) | 0.0088<br>(0.0041–0.0154) | 0.0081<br>(0.0021–0.0172) | 0.0029<br>(0.0003–0.0077) | 0.0051<br>(0.0012–0.011) | 0.0047<br>(0.0007–0.0109) | 0.0119<br>(0.0034–0.0232) |
| $\hat{\pi}_{int}$ | 0.0034<br>(0.0005–0.0097) | 0.0049<br>(0.0017–0.0099) | 0.0048<br>(0.0006–0.0121) | 0.0018<br>(0–0.0055) | 0.0028<br>(0–0.0077) | 0.0027<br>(0–0.0074) | 0.0067<br>(0.0014–0.0148) |
| $\hat{\theta}_n$ | 0.0012<br>(0.0003–0.0027) | 0.0015<br>(0.0005–0.0029) | 0.0011<br>(0.0002–0.0023) | 0.0005<br>(0–0.0012) | 0.0006<br>(0–0.0013) | 0.0005<br>(0–0.0013) | 0.0011<br>(0.0002–0.0026) |
| $\hat{\theta}_s$ | 0.0073<br>(0.0022–0.0166) | 0.0098<br>(0.0049–0.0166) | 0.0079<br>(0.0023–0.0161) | 0.0027<br>(0.0003–0.0067) | 0.0044<br>(0.001–0.0094) | 0.0038<br>(0.0006–0.0087) | 0.0096<br>(0.0029–0.019) |
| $\hat{\theta}_{int}$ | 0.0042<br>(0.0009–0.0106) | 0.0054<br>(0.0021–0.0099) | 0.0046<br>(0.0008–0.0111) | 0.0016<br>(0–0.0044) | 0.0024<br>(0–0.0065) | 0.0022<br>(0–0.006) | 0.0054<br>(0.0012–0.0118) |
| *H<sub>n</sub>* | -0.1198<br>(-0.6092–0.1929) | -0.0635<br>(-0.2935–0.1128) | -0.0274<br>(-0.1759–0.1143) | -0.0589<br>(-0.3115–0.9992) | 0.0137<br>(-0.1256–1.2172) | -0.0114<br>(-0.2371–1.2172) | 0.0246<br>(-0.078–0.5348) |
| *H<sub>s</sub>* | -2.2802<br>(-5.4425–1.1004) | -0.1197<br>(-0.3405–0.0735) | -0.0444<br>(-0.1897–0.0858) | -0.1074<br>(-0.2942–0.9992) | 0.0006<br>(-0.131–0.0789) | -0.02<br>(-0.1177–0.0774) | -0.0163<br>(-0.0539–0.0326) |
| *K<sub>n</sub>* | 0.0215<br>(0.0077–0.0415) | 0.0203<br>(0.007–0.0404) | 0.0216<br>(0.0079–0.04) | 0.0221<br>(0.0081–0.0404) | 0.0201<br>(0.0076–0.0378) | 0.0213<br>(0.008–0.039) | 0.0214<br>(0.0082–0.0402) |
| *K<sub>s</sub>* | 0.1394<br>(0.0884–0.1876) | 0.1369<br>(0.0892–0.1887) | 0.1439<br>(0.0927–0.1878) | 0.1533<br>(0.103–0.2056) | 0.1508<br>(0.1011–0.2037) | 0.1624<br>(0.1075–0.2168) | 0.1541<br>(0.1009–0.203) |
| NI | 1.1798<br>(0.3653–2.6753) | 1.185<br>(0.4161–2.2767) | 1.0788<br>(0.2228–2.6991) | 1.614<br>(0–4.6016) | 1.3401<br>(0–3.9734) | 1.271<br>(0–3.5996) | 0.9777<br>(0.2139–2.2241) |
| CAI | 0.419<br>(0.2787–0.5453) | 0.4167<br>(0.2794–0.5331) | 0.4138<br>(0.2725–0.5288) | 0.4045<br>(0.2619–0.5255) | 0.3973<br>(0.2674–0.5242) | 0.3991<br>(0.2577–0.5174) | 0.4105<br>(0.2641–0.5274) |

The MKT is susceptible to violations of the standard neutral model. In particular, demographic change and the segregation of slightly deleterious mutations can bias *α* estimates unworldly or downwardly (Eyre-Walker 2006; Messer and Petrov 2013a). One way that we accounted for this bias was by excluding low-frequency variants in the standard MKT. The other way was by calculating *α* and *ω* using the distribution of fitness effects (DFE: *α*_*dfe*_ and *ω*_*α*_) (Eyre-Walker and Keightley 2007, 2009). Notably, the proportion of adaptive fixations, *α*_*dfe*_, resulted in higher estimations based on the standard MKT test (Fig 3A). We also note higher between species variation, both with respect to *α*_*dfe*_ and (Fig. 3). Mean *α*_*dfe*_ values ranged from 63% in *A. jacksonii* to 11% in *A.* sp-jack1 (Fig. 3A). Given that results based on the DFE indicate an underestimation of *α* it would be conservative to consider genes with *α* > 0 under positive selection. The number of genes with at least one adaptive mutation per 1,000 bp (i.e. *α* > 0.1%) were 286 in *A.* sp-F11, 280 in *A.* sp-jack2, 240 in *A. jacksonii*, 216 in *A.* sp-jack5/6, 213 in *A.* sp-jack1, 203 in *A.* sp-T31, and 151 in *A.* sp-jack3. Besides *α*_*dfe*_ and *ω*_*α*_, the DFE can also be used to estimate the proportions of deleterious mutations under different selection strengths (scaled by the effective population size, *N_e_s*). The DFE estimates up to four different categories, namely *N_e_s* < 1, 1 < *N_e_s* < 10, 10 < *N_e_s* < 100 and *N_e_s* > 100 Most species had a higher proportion of mutations with very strong effects (Fig. 4; e.g. *N_e_s* > 100), with proportions between 0.4 and 0.8 (except *A.* sp-T31, which had a median lower than 0.4). *Amanita jacksonii* and *A.* sp-T31 had the highest proportion of mutations with intermediate effects (10 < *N_e_s* < 100), while categories with *N_e_s* < 10 had proportions of less than 0.2 for all species (Fig. 4).

**Figure 3.**
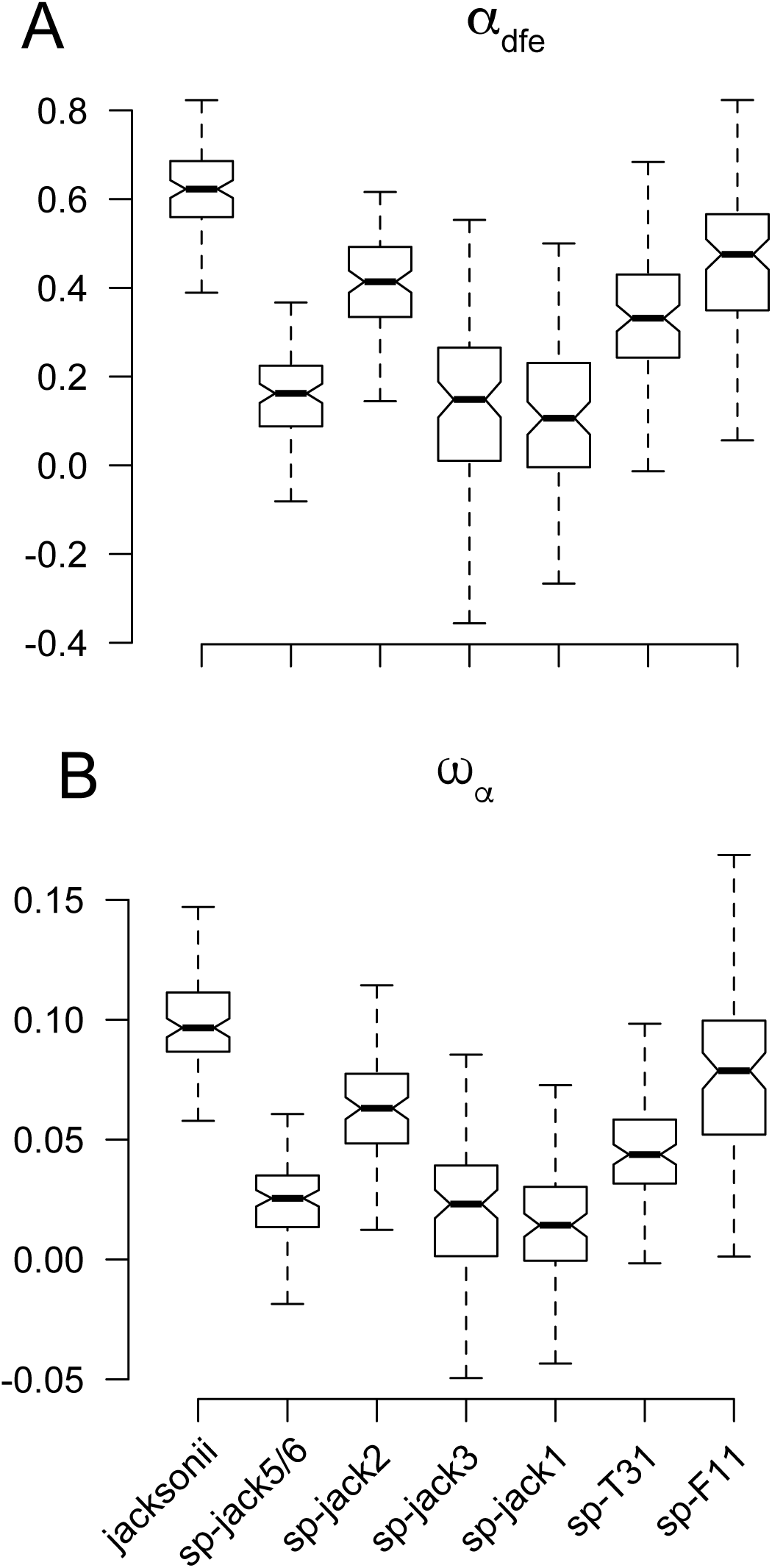
Estimates of (A) the proportion of adaptive substitutions (α_dfe_) and (B) the rate of adaptive evolution (*ω*_α_) based on the distribution of fitness effects.

**Figure 4.**
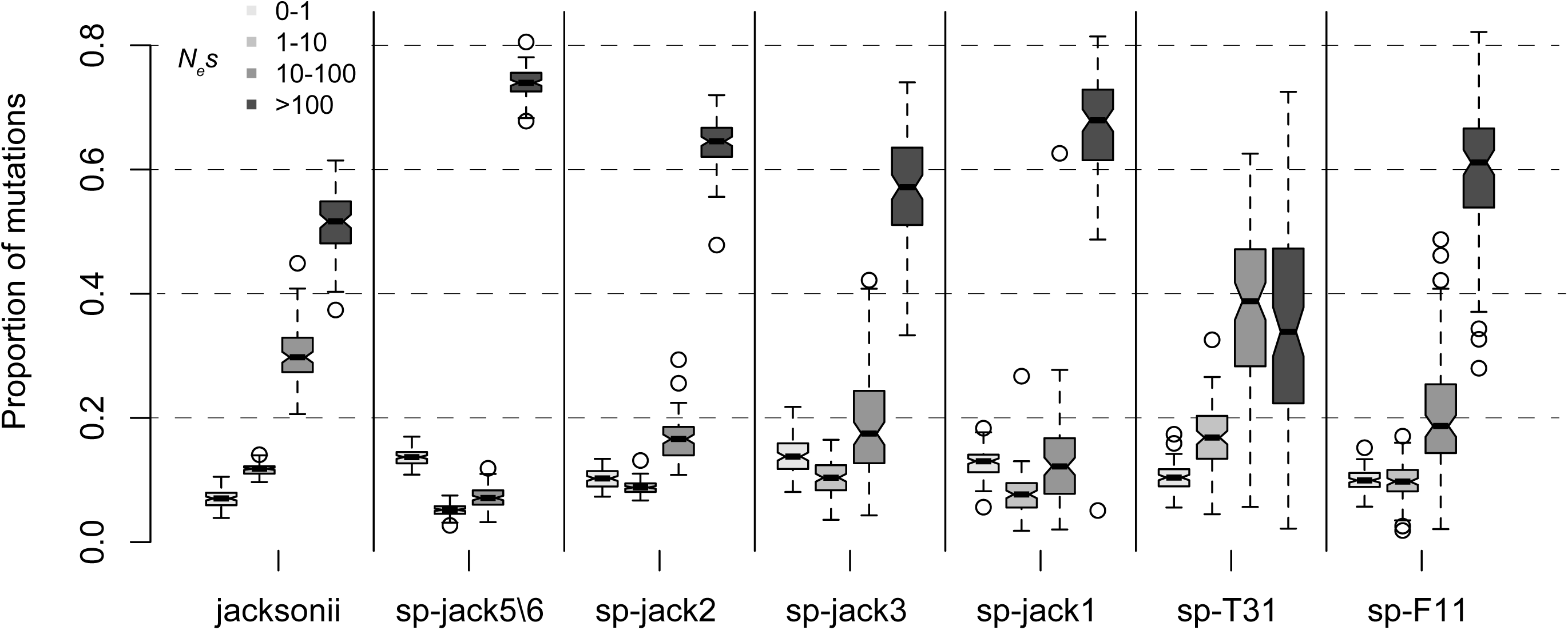
The proportion of deleterious mutations under four different *N_e^s^_* categories based on the distribution of fitness effects model.

### Historical population sizes

We inferred the demographic history of each species using extended Bayesian skyline model (eBSP, Heled and Drummond 2008) implemented in BEAST v1.8.2 (Drummond et al. 2012). We opted for reducing our data set in demographic history estimation for several reasons: (1) it has been shown that 16 informative genes or more should be sufficient to estimate accurate population size histories (Heled and Drummond 2008); (2) some of the genes we sampled are contiguous and possibly linked, biasing the assumption of free recombination between loci; and (3) to improve Bayesian parameter mixing and convergence times, which increases with more data (Suchard and Rambaut 2009).

The eBSP analyses supported at least one demographic change within the 95% posterior density distribution (PDD) in all seven species. The species with the highest mean number of demographic changes was *A.* sp-F11 with 1.37 [1, 2 95% PDD], while the lowest was in *A.* sp-jack1 with 0.3 [0, 1 95% PDD]. Demographic trends of population size expansions were observed in *A. jacksonii, A.* sp-jack5/6, *A.* sp-jack2, and *A.* sp-F11, and to some extent in *A.* sp-T31, while *A.* sp-jack3 and *A.* sp-jack1 had a rather constant demographic trend (Fig. 5). The most abrupt demographic expansions were found in *A. jacksonii* and *A.* sp-jack5/6 with a near 6 to 7-fold increase in population size within the last ca. 2 Myr (Fig. 5). Similar estimates were found in the two-phase demographic model in the DFE analysis.

**Figure 5.**
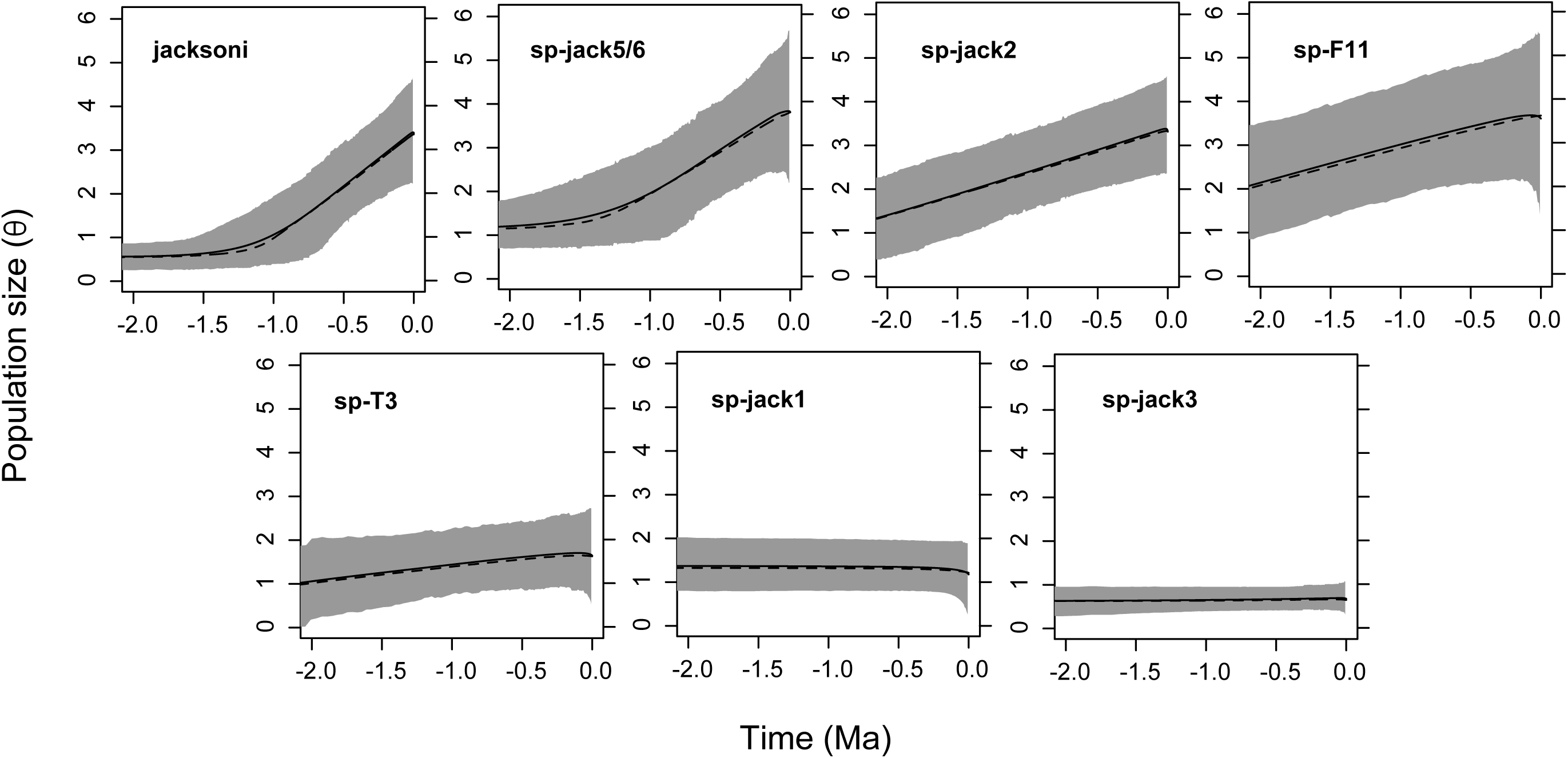
Demographic trends through time for each species based on the eBSP model. Grey shading represents 95% posterior confidence intervals (CI). Solid lines represent mean population size values and dotted lines represent median values.

### Gene Ontology

Due to functional constrain, single-copy genes typically fall within specific functional groups, for instance, in binding and catalytic activities (Aguileta et al. 2008; Han et al. 2014). We find a similar functional bias (Fig. 6A), where a large proportion included proteins involved with energetic and metabolic processes (ATP binding, GO:0005524) and protein-protein interactions (protein binding, GO:0005515, Fig. 6). To less extent proteins related directly or indirectly with gene expression regulation (i.e. nucleotide [GO:0000166], nucleic acid [GO:0003676], and DNA [GO:0003677] binding; zinc ion binding [GO:0008270]; regulation of transcription, DNA-dependent [GO:0006355]), transport and signaling (i.e. protein phosphorylation [GO:0006468], phosphorylation [GO:0016310], protein kinase activity [GO:0004672, GO:0004712], trans-membrane transport [GO:0055085], protein transport [GO:0015031]), metabolic processes (i.e. oxidation-reduction processes [GO:0055114], oxidoreductase activity [GO:0016491], carbohydrate metabolic process [GO:0005975], hydrolase activity [GO:0016787]), and homeostasis (calcium ion binding [GO:0005509]) were also found in the mix. Nonetheless, we accounted for overrepresented Gene Ontology (GO) terms by adding up the positive selection signal (e.g. *α* and the proportion of sites with *ω* > 1) across genes and species and then scaling by the total number of unique GO terms (see Materials and Methods). Some of the GO terms found in genes under strong directional selection included proteins related to transcription activities, cellular transport, and metabolic activities (Fig. 6B). Interestingly, GO term composition of genes under diversifying selection varied from that of genes under directional selection (e.g.*α* > 0), finding stronger selection in proteins with functions related to biosynthetic and other metabolic processes (Fig. 6b). From the 474 genes across all species were found with, *α* > 0, 17 were found exclusively in *A.* sp-F11, followed by 10 in *A.* sp-T31 and sp_jack5/6, 8 in *A. jacksonii* and *A.* sp-jack2, 7 in *A.* sp-jack1, 6 in *A.* sp-jack5/6. Information about their GO and annotation can be found in Table S1 Online Supplementary Data.

**Figure 6.**
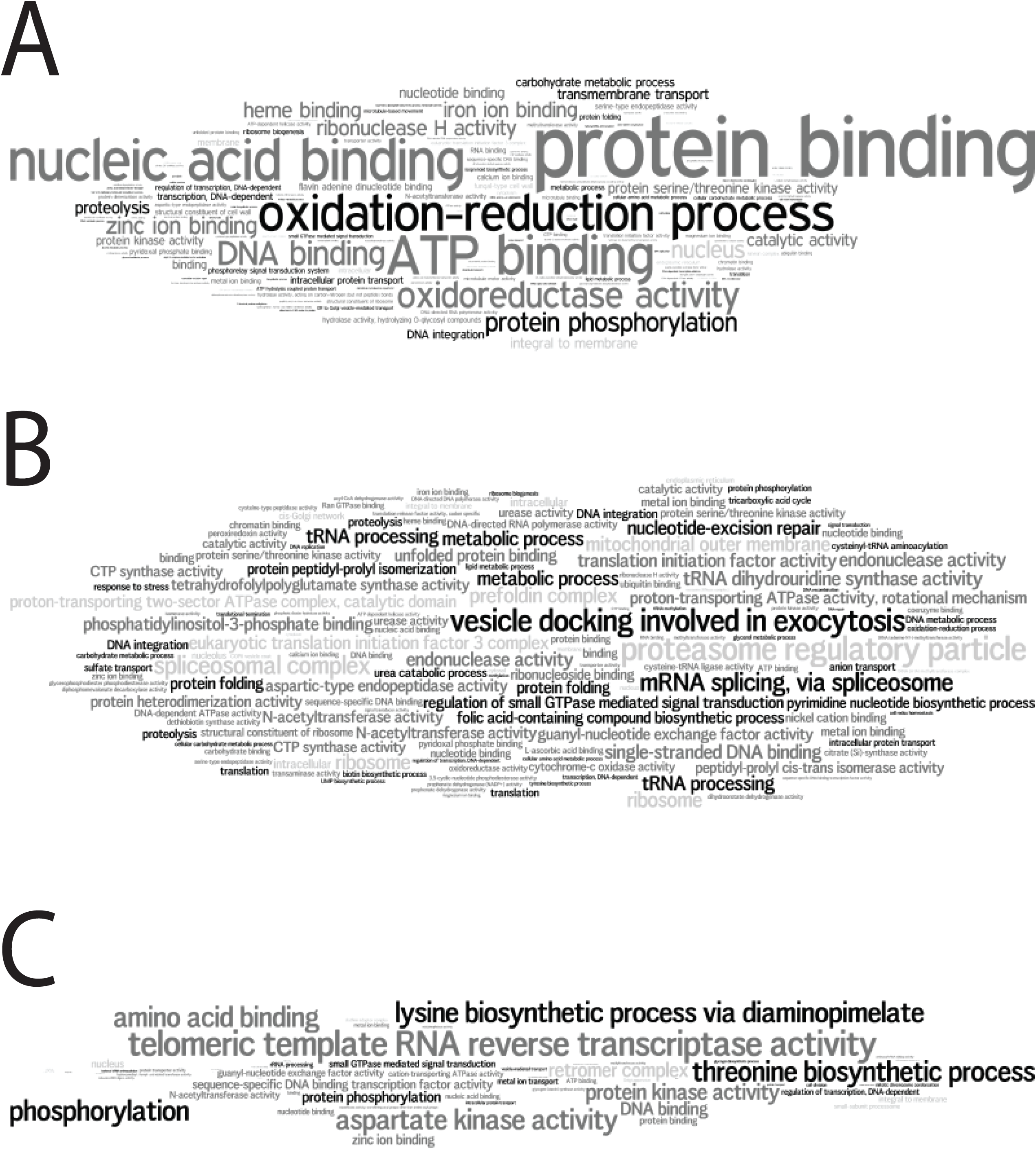
Gene Ontology terms *Wordles* for (A) all genes in the data set, (B) genes with α > 1, and genes with a significant proportion of sites under diversifying selection. The size of the term is equivalent to the extent of selection and how often the term was found across genes and species.

## Discussion

Interpreting DNA variation from an evolutionary perspective has been a long-standing goal for population geneticists, both doing empirical and theoretical work. The neutral equilibrium theory of molecular evolution (Kimura 1977, 1986) provides a null framework to test hypotheses about natural selection and other processes acting on the genome. While the core of population genomic research has developed around coalescent-based theory in a single –or at least a pair– of species (Hough et al. 2013; Charlesworth 2010), there are clear benefits in integrating comparative and phylogenetic data, in multi-species multi-locus assessments (Cutter 2013).

### The exon-targeted sequence-capture approach

Before and early into the “whole-genome sequencing” era, most population genomics studies relied on PCR and Sanger sequencing of individual genes for data production. Some classic studies included data ranging from tens to few hundreds of genes, in what were then called “whole-genome” approaches. Exon-targeted sequencing takes advantage of the rapid and massive data production of next-generation sequencing, while focusing on specific genes of interest. Although any class of gene can be potentially targeted, we opted for single-copy genes, attempting to avoid potential spurious DNA hybridizations with paralogues, which can bias diversity estimates. However, targeting single-copy genes has the disadvantage of characterizing genome-wide diversity based on a single “class” of genes, which may not necessarily represent a heterogeneous sample from the genome. Nonetheless, the degree of variation and distribution of the data within each species, in combination with our results, suggests that our sample of 502 genes maybe a good representation genome-wide variation. Furthermore, this method has been proven successful when DNA material is only available in low quantities (Hancock-Hanser et al. 2013), or from degraded samples (Templeton et al. 2013). All of our samples came from dried museum specimens and FTA plant saver cards (Dentinger et al. 2010), which would not reach quantity/quality standards for other popular downstream genomic applications such as whole-genome resequencing or reduced-library representation approaches (e.g. RAD-tag, GBS). In particular, reduced-library representation approaches are practical for sampling genome-wide SNPs, however the shortness of the fragments makes it impossible to conduct codon-based divergence and polymorphism assessments. Therefore, exon-targeting methods are a practical and efficient strategy for scoring many loci and complete structural genes in non-model organisms.

### Effects of selection and demography on genomic variation

Studies in flies (Smith and Eyre-Walker 2002; Bierne and Eyre-Walker 2004; Sella et al. 2009), plants (Slotte et al. 2010), and humans (Fay et al. 2001) have suggested that the rate of adaptive substitution can reach high proportions (20–40%), even in the presence of high levels of selective constrain or purifying selection (Williamson et al. 2014; Slotte et al. 2010). The rate of adaptive evolution is thought to be even higher in microorganisms, such as bacteria (Charlesworth and Eyre-Walker 2006). In contrast, in these *Amanita* species, *α* seems to be consistently underestimated (Fig. 2C). A number of different factors can explain potentially low levels of adaptive evolution. Among them, we will discuss how specific evolutionary scenarios such as purifying selection, relaxed selective constrain, and demographic history can contribute to this pattern, highlighting evidence in favor or against such factors.

Strong selective constraint is expected in protein coding DNA, largely because the majority of amino acid changing mutations are likely to be deleterious (Ohta 1992; Andolfatto 2005). One possible explanation for values of *α* ~ 0 is purifying selection. If non-synonymous substitutions are not being fixed between species, it is likely that they were deleterious early in the evolution of a lineage, and thus purged from the population (Fay and Wu 2003), typically leading to a low ratio of 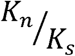 Both polymorphism (Table 1) and divergence (Fig. 2B) data overall support high selective constraint in the *A. jacksonii* complex. Similarly, codon LRTs indicate that 90% of the genes favor a model in which the majority of sites are under long-term purifying selection. Most species also resulted with high proportions of new (deleterious) mutations under strong selective effects (e.g. *N_e_s* > 10; Fig. 4). We believe that this strong signal of high purifying selection could, in part, be a reflection of a functional bias in single-copy genes (Fig. 6). Many single-copy genes are thought to be under strong selective constrain for duplication, which is also linked to functional constrain (De Smet et al. 2013). Other studies in plants and fungi have characterized single-copy genes under conserved cellular functions, such as binding and metabolism (Aguileta et al. 2008; Han et al. 2014).

Moreover, mixed genome-wide signals of positive and negative selection are not uncommon (Williamson et al. 2014; Fay et al. 2001), and our results also show evidence of positive selection. While we find little support for extensive diversifying selection, as only 10% of genes supported model M2a (site class with *ω* > 1), more genes were found under directional selection within and across species. On average each species had about half the total number of genes (ca. 230 from 502) with *α* > 0, and about 94% (474) of all 502 genes had at least one species with *α* > 0. However, only 25 genes where found with *α* > 0 for all seven species, which is a number closer to the number of genes found under diversifying selection. The diversifying selection model requires selection in favor of allelic diversity, which may rarely happen in genes under functional constraint (Yang et al. 2000). Nonetheless, molecular adaptation is possible, even when 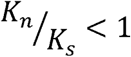(Kryazhimskiy and Plotkin 2008).

The fate of adaptive and slightly deleterious mutations is highly affected by *N*_*e*_ (Ohta 1992; Eyre-Walker and Keightley 2007; Charlesworth 2009). Standard MKT approaches are generally insensitive to the presence of slightly deleterious mutations and/or demographic changes, which can bias *α* estimates (Smith and Eyre-Walker 2002; Eyre-Walker and Keightley 2009; Messer and Petrov 2013a). Methods such as the DFE model can both account for slightly deleterious (non-synonymous) mutations that tend to contribute more to polymorphism than to divergence, as well as demographic changes (Eyre-Walker and Keightley 2007, 2009). Our results show that *α* (standard MKT) is largely underestimated across all species (Fig. 3). This is more apparent in species that have undergone recent population expansions (Fig. 5). Slightly deleterious polymorphisms are more likely to result in underestimation of *α* in species with low *N_e_* or that have undergone a population contraction (Smith and Eyre-Walker 2002; Charlesworth 2009; Eyre-Walker and Keightley 2009). We note that effectively neutral non-synonymous mutations with *N_e_s* ≈ 1 are less than 20% in all species (Fig. 4), however, *α* (standard MKT) is strongly underestimated in species with historically low *N_e_* (Fig. 5).

Untangling the effects of demography and selection on patterns of polymorphism is difficult (Andolfatto 2001). In particular, because certain demographic scenarios, such as population expansions, can result in similar polymorphism footprints. Nevertheless, demographic events are more likely to affect the whole genome rather than individual loci in the case of selection (Stajich and Hahn 2005). We found marked differences in Tajima’s *D* between species indicating different levels of skewness in the site frequency spectrum (SFS), in particular, the negative skew found in *A. jacksonii* (Fig. 2a). In this species, a previous study has evidenced an increase in population size linked to northward population expansion (Sánchez-Ramírez et al. 2015b). Here we corroborate with a larger genomic sample that this expansion signature (Fig. 5) is likely to be legitimate, but also note that it may be exacerbated by the effects of positive selection (Fig. 3). Overall, specific patterns of demographic history are consistent with Tajima’s *D* values. For instance, species with positive Tajima’s *D* values tended to have more constant and smaller population sizes. In comparison, species with negative *D* had expansion signatures and larger population sizes (Fig. 2A, 5). Sánchez-Ramírez et al. (2015b) found expansion signatures in *A.* sp-jack2 and *A.* sp-jack5/6, which are also supported by our results at a higher genomic scale (Fig. 5). Similarly, we find consistent demographic scenarios in *A.* sp-jack3, *A.* sp-jack1, and *A.* sp-T31, as reported in Sánchez-Ramírez et al. (2015b). We find particularly interesting contrasting demographic and molecular evolution patterns in two species: *A. jacksonii* and *A.* sp-F11. On one side, they have opposite site frequency skews. While *A. jacksonii* has tendency for low and high frequency variants (a negative *D* and *H*; Fig. 2a and Table 1), *A.* sp-F11 only has a tendency for intermediate frequency variants (positive *D*; Fig. 2a). This alone could be explained by differences in their demographic history (Fig. 5), where the former has suffered a bottleneck previous to a dramatic expansion northward, while the latter has had a more stable range and demography, with comparatively high *N_e_* (Fig. 1 and 5). While these differences are clear, both species have the highest *α*_dfe_ values in the complex (Fig. 3), suggesting high proportions of adaptive evolution. One possibility for this apparent contradiction is that, in *A. jacksonii*, population size expansion could have led to the fixation of slightly deleterious mutations, contributing disproportionally to divergence and overestimating *α*_dfe_ (Eyre-Walker and Keightley 2009). Another explanation is that glacial cycles during the Pleistocene could have acted as fluctuating episodes of selection, which can lead to high levels of adaptive evolution (Huerta-Sanchez et al. 2008; Bell 2009; Gossmann et al. 2014). This group most likely diversified within the last 5 Myr (Fig. 1B; Sánchez-Ramírez et al. 2015b), during a period that is characterized by a series of abrupt temperature oscillations that have deeply affected the genetic diversity of many populations and species (Hewitt 1996, 2000, 2004; Hoffmann and Sgrò 2011)

### Functional bias, adaptive genes, and the EM habit

The genes that were assayed here were not selected for their specific functions. In fact, they were selected due to their single-copy nature, which facilities bait homology and specificity in exon-targeting procedures, and lowers the rate of alignment artifacts. Nonetheless, single-copy genes are known to be functionally constrained (De Smet et al. 2013), which can introduce noise when assessing functional enrichments of genes under selection. In order to reduce the effect of overrepresentation of GO term due to this functional bias, we devised a measure to scale down the amount of selection in overrepresented genes (see Material and Methods). Differences in abundance of GO terms in the overall data set (Fig. 6A) and GO terms found in genes under selection (Fig. 6A, B), suggest that the overrepresentation was mitigated. Overall, protein functions of genes under selection seem to be divided into three general categories; those related to (1) gene transcription/expression activities, (2) signalling and transport, and (3) metabolic oxidative processes (Fig. 6). Many of these functions coincide with divergent regions in the genome of another EM fungus (*S. brevipes*) that are also thought to be under selection (Branco et al. 2015). We also noted some functions related to carbohydrate metabolism, specifically with catalytic and hydrolytic activities (Fig. 6A, B). These may be compelling because all EM fungi depend on external carbohydrate sources that are generally supplied by their hosts (Nehls et al. 2007). Throughout the evolution of the EM symbiosis, many species have suffered from a convergent loss of decay mechanisms, while retaining moderate gene repertoires with lignocellulitic capabilities (Kohler et al. 2015). Adaptation in proteins related to carbohydrate metabolism, such as hydrolase activity, could allow for a more efficient breakdown of complex polysaccharides during periods of stress and/or low photosynthetic activity. Also worth mentioning, proteins with telomeric template reverse transcriptase activity were found under diversifying selection (Fig. 6C). Studies suggest that the dynamic nature of telomeric and subtelomeric regions are likely to contribute to rapid evolution of fungi during biotic interactions (Aguileta et al. 2009).

## Material and Methods

### Samples and whole-genome data

Sixty-three dried specimens, representing eight species (Sánchez-Ramírez et al. 2015b) were processed from collections made in various locations in North America, all deposited in fungal herbaria (Table S2). DNA was extracted from ca. 30-50 mg of dried fungal tissue (gill) and extracted using a modified standard CTAB/proteinase K/chorophrom:isoamilic alcohol protocol (van der Nest et al. 2014). In addition, whole-genome 454 sequences were produced by the Duke Center for Genomic and Computational Biology for a North American species outside the *A. jacksonii* complex –*A. basii*. These sequences were mapped onto the *A. jacksonii* TRTC168611 draft genome (van der Nest et al. 2014), with the purpose of building a whole-genome draft assembly of an outgroup species.

Identification of single-copy orthologs, probe design, and gene selection

Structural genes were predicted in *A. jacksonii* and *A. basii* using AUGUSTUS v3.0.3 (Stanke et al. 2006). All CDS in each species were reciprocally aligned against each other using command-line (standalone) BLASTn, creating two files in tabular format (-outfmt 7). Next, a custom perl script (rbh.pl, available from: https://sites.google.com/site/santiagosnchezrmirez/home/software/perl) was used to find reciprocal single-copy hits, putting each pair in a single FASTA file. Each single-copy pair was aligned using MUSCLE v3.6 (Edgar 2004). Probes for target hybridization were designed based on conserved (identical) CDS regions, 60bp long, across both species. Only genes with four or more non-overlapping regions were chosen. Two PCR primers (5’-TAATACGACTCACTATAGGG-3’ and 5’-CTATAGTGTCACCTAAATC-3’) were added to the 5’ and 3’ ends of the 60bp probe target, for a total of 100bp per oligonucleotide probe.

### Sequence capture, library preparation, and sequencing

Custom oligos probes were synthesized by LCScience (http://www.lcsciences.com/applications/genomics/oligomix/) using DNA microchip technology (Gao et al. 2004). The synthesized probes were then PCR amplified as needed for downstream applications. Probes were later transcribed into RNA using MegaScript T7 Kit (Invitrogen, Carlsbad, CA) and Biotin-dUTP to produce labeled oligos. The DNA was sheared and a standard Illumina TruSeq V2 DNA library kit was prepared, multiplexing all 46 samples. Sequence captures (probe-target hybridization) were performed with 200ng of library, 500ng of biotinylated probes, and Steptavidin beads (Invitrogen, Dynabeads M-270, CAT65305). The capture procedure was repeated at least twice. The details of the protocols can be found in Supplementary file 1. All target captures were directly sequenced in a single Illumina Hi-Seq 2500 lane.

### Bioinformatics

#### *Read alignments, sequence phasing*, and *quality control*

Sequences were de-multiplexed into individual fastQ files and each one mapped onto the targets’ whole-gene sequences from *A. jacksonii* using the Burrows-Wheeler Aligner’s (BWA v0.7.12) algorithm BWA-MEM (Li and Durbin 2010). The resulting SAM files were then filtered with *view* in SAMtools v0.1.19 (Li et al. 2009) for off target reads, low quality mapping (< 30), and PCR duplicates, and converted to BAM files. The reference sequences were indexed and the BAM files sorted. We used HapCompass (Aguiar and Istrail 2012), which uses a read-based graph algorithm, to generate haploid sequences from BAM and VCF files. The VCF file was produced by “piping” *mpileup*, bcftools’ *view*, calling the “-cg” flags for genotype likelihood computing, and *varFilter* in vcfutils.pl, only keeping SNPs with Fred quality ≥ 20. The SAM files produced by HapCompass were then converted to BAM and sorted. Consensus fastQ files were produced by “piping” *mpileup*, bcftools’ *view*, and *vcf2fq* in vcfutils.pl, in a similar way as above. A custom perl script (OrderFromSamtools.pl) was used to convert fastQ files to fastA format and to produce by-gene alignments from by-sample files. A second perl script (getPhasedGenome.pl) parsed those fastA files to produce “a” and “b” allelic variants per haploid sequence. All missing data was scored as Ns, while unsorted polymorphisms were either left as IUPAC ambiguity codes or scored as Ns, depending on the analysis.

#### Phylogenomics and lineage sorting

We used MrBayes v3.2 (Ronquist et al. 2012) to generate individual gene trees based on the general time reversible (GTR) model and among-site rate heterogeneity modeled as a gamma distribution (+G). For each gene, we ran two parallel runs each with eight Metropolis-coupled Markov Chain Monte Carlo (MC^3^) chains and 1 million generations. We sampled every 100^th^ state and discarded the 10% initial states as burnin. We assessed convergence by making sure that the average standard deviation of split frequencies was ≤ 0.01. Posterior trees were summarized onto a majority-rule consensus tree compatible with clades with frequencies ≥ 0.5. The summary trees were imported into R using *ape* (Paradis et al. 2004) where the genealogical sorting index (*gsi*, Cummings et al. 2008) was calculated. The *gsi* measures, in a range from 0 to 1, the degree of exclusive ancestry in labeled terminal groups, where 1 signifies group monophyly. These groups can represent any type of biological association, such as species or populations.

### Analyses

#### DNA polymorphism and divergence

Within species all sites were bi-allelic. Coding regions were based on the annotation of *A. jacksonii* (van der Nest et al. 2014). In coding regions, sites were separated into synonymous (4-fold, 3-fold, and 2-fold degenerate sites) and nonsynonymous (0-fold degenerate sites) using the method of Nei and Gojobori (1986). Polymorphism was measured as the average number of pairwise nucleotide differences per site between any two random DNA sequences or 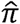 (Nei and Li 1979; Nei 1987), or the Watterson’s estimator 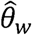, which is measured as the proportion of segregating sites *S* in a sequence of length *L* divided by the (*n* − 1) th harmonic mean number. Both 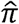 and 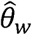 were estimated using SFS counts based on formulae in Campos et al. (2014). To measure the skewness of SFS we calculated Tajima’s *D* (Tajima 1989) and Fay and Wu’s *H*, which measures skewness in the derived frequency spectra following Zeng et al. (2006). We measure codon usage bias by calculating the codon adaptation index (CAI, Sharp and Li 1987). Divergence *K* was measured as the Kimura-2-parameter (Kimura 1980) proportion of sites fixed in the ingroup compared to the outgroup (*A. basii*), considering transitions and transversions separately. In order to control for low-frequency ascertainment biases, we excluded segregating sites with a frequency of 1/N within species, where N is the total number of sequences per species. All calculations were performed using the script *poly+div_sfs.pl*, available at https://sites.google.com/site/santiagosnchezrmirez/home/software/perl. For comparison, estimations were also computed in DnaSP v5 (Librado and Rozas 2009) and Polymorphorama (Bachtrog and Andolfatto 2006, http://ib.berkeley.edu/labs/bachtrog/data/polyMORPHOrama/polyMORPHOrama.html).

#### McDonald-Kreitman tests

We estimated the degree of adaptive evolution in CDS for each species by measuring the amount of polymorphism (within species DNA changes) and divergence (fixed DNA changes between species) leading to non-synonymous and synonymous substitutions. We used the McDonald-Kreitman test (MKT) (McDonald and Kreitman 1991) to estimate the Neutrality Index (NI),

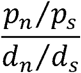

Where *p_n_* and *p_s_* are, respectively, the within-species proportion of non-synonymous and synonymous segregating sites per non-synonymous and synonymous site, and *d_n_* and *d_s_*, the proportion of fixed non-synonymous and synonymous substitutions per non-synonymous and synonymous site. Values closer to 1 indicate neutrality, those close to 0 indicate positive selection, and those > 1 suggest either negative purifying selection or balancing selection. We used the formula derived by (Smith and Eyre-Walker 2002) to estimate the proportion of amino acid substitutions under positive selection:

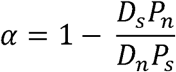

This model assumes that mutations are either strongly deleterious or neutral, thus model violations can arise in the presence of slightly deleterious mutations. Likewise, demographic changes can also cause *α* to be under or overestimated (Eyre-Walker 2006). In order, to account for demography and slightly deleterious mutations we used the approach by Eyre-Walker and Keightley (2007, 2009), which uses a maximum likelihood approximation based on the distribution of fitness effects (Keightley and Eyre-Walker 2010; http://www.homepages.ed.ac.uk/pkeightl//dfe_alpha/dfe-alpha-download.html). Here, estimated the derived SFS for each gene and for each species by polarizing alleles using the outgroup (*A. basii*). We first used *est_dfe* on the folded “neutral” and “selected” (e.g. synonymous and non-synonymous sites) SFS to estimate the DFE based on a two-stage demographic model fixing the *N*_1_ to 100 and optimizing *N*_2_. The mean selection coefficient *s* and the *β* parameter were also optimized using as starting values −0.1 and 0.5 respectively. We then ran *est_alpha_omega* specifying “neutral” and “selected” counts of diverged (fixed) sites. We applied Jukes-Cantor correction for multiple hits when estimating divergence and removed polymorphism contributing to divergence. In order to generate confidence intervals, we re-estimated parameters by bootstrapping 100 times 100 randomly sampled SFS. Every time the 100 SFS were summarized for each allelic bin.

#### Codon-based model selection

Models that solely rely on *K_n_/K_s_* (*ω*) (ratios are also robust and popular methods for detecting natural selection at the molecular level (Nielsen 2001; Yang and Nielsen 2002). Essentially, CDS regions that have an equal number non-synonymous and synonymous mutations are thought to evolve under effective neutrality, having a *ω* = 1 Deviations from this ratio indicate purifying selection if *ω* < 1 and diversifying (positive) selection if *ω* > 1 (Yang 1998; Anisimova et al. 2001). We used likelihood ratio tests (LRT) to discriminate between three codon-based models: (1) a site-wise model with a single *ω* ratio; (2) a site-wise “nearly neutral” model where there is a proportion of sites *p_0_* with, *ω*_0_ < 1 and a proportion of sites *p*_0_ with *ω*_1_ = 1 This serves as a null model for a third that considers an additional category of sites with *p*_2_ with *ω*_2_ > 1 In the last two models, *p*_1_ and *p*_2_ are fixed as *p*_1_ = 1 – *p*_0_ and *p*_2_ = 1 – *p*_0_ – *p*_1_, respectively. All models are based on Goldman and Yang’s (1994) codon substitution model and were estimated using maximum likelihood in PAML v4.8a (Yang 2007) using *codeml*. Because *ω* ratios assume that mutations are fixed differences between species and because population polymorphisms may potentially bias *ω* estimates (Kryazhimskiy and Plotkin 2008), we selected only one sequence per species per gene.

#### Historical demography

To infer the demographic history of each species, we applied the multi-locus eBSP model (Heled and Drummond 2008) using the Kimura-2-parameter (K80) model with gamma distribution as the substitution model, implemented in BEAST v1.8.2 (Drummond et al. 2012). As data, we selected 29 intron regions that complied to a series of filters, namely a length size of at least 400 bp and locations in different linkage groups. This last filter was done to ensure that loci are not in linkage disequilibrium, as the eBSP model assumes free between loci recombination. In each species, we included an alignment the translation elongation factor 1 alpha (*tef1*), which has been used as molecular clock proxy with a substitution rate of 0.00194 substitution/site/Myr (Sánchez-Ramírez et al. 2015b). The substitution rate in the other genes is scaled based on the fixed rate. Each of the eBSP analyses ran for 100 million generations, with a sampling rate of 10%, and a burnin of 10%. We assessed convergence and mixing by looking at likelihood per generation trace plots and effective sample size values in Tracer 1.6 (Rambaut et al. 2013). Population size values in eBSP were produced directly from the analyses in CSV format.

#### Structural and functional annotations

Whole-gene sequences from *A. jacksonii* were translated into amino acids and then imported into Blast2GO (Conesa et al. 2005), which was used as an annotation tool. The main amino acid sequence annotation was taken from the non-redundant (nr) protein database in NCBI using BLASTp, only keeping the best 10 matches. In addition, we ran InterProScan v5 (Zdobnov and Apweiler 2001) to identify functional domains, from which we extracted GO terms. For visualization purposes, we used advanced “word clouds” in http://www.wordle.net/advanced, where scaled GO terms were laid out. Molecular function, biological processes, and cellular component, were marked by different colors. For the scaling proportion, we used:

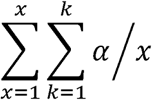

where *α* is the value of the amount of adaptive evolution (*α* or *P*_*ω*_) summed over *k* species which have found *α* > 0 and summed over *x* times a given GO term was found. In the case of *α k* and *x* can be minimaly 1, however for *P*_*ω*_, which is the proportion of sites with *ω* < 1, there is only a single value per gene (i.e. *k* always equals 1), but *x* can be > 1.

## Acknowledgements

We would like to thank Hernán López-Fernández and Katriina Ilves for sharing their experiences with exon-targeting sequencing, and Dax Torti for providing advice on best library preparation and sequencing practices. We also acknowledge SciNet and CAGEF for providing access to high-performance computing infrastructure. Funding was provided through grants to J.M.M. by the Royal Ontario Museum (ROM) Governors and Natural Science and Engineering Research Council (NSERC) of Canada, and a scholarship by the National Science and Technology Council of Mexico (CONACYT) for doctoral studies to S.S.R.

## Supplementary data

Table S1. GO terms of singleton genes with α > 1.

Table S2. Species, specimen voucher name, and geographic location data for all samples.

Figure S1. Variable sites and percentage of missing data plotted against gene lenght.

Figure S2. Sample of 100 genealogies from 502 estimated trees. Terminal colors indicate different species.

Figure S3. Distribution of *gsi* values for all genes and species. Species with higher amounts of conflict are highlight.

Figure S4. Scatterplot of Tajima’s *D* in synonymous and non-synonymous sites against the *gsi*. Grey points highlight reciprocally monophyletic genes.

Figure S5. Scatterplot of α values against the *gsi*.

Figure S6. Scatterplot of α values and synonymous nucleotide diversity against codon usage bias (CAI).

